# Molnupiravir (MK-4482) is efficacious against Omicron and other SARS-CoV-2 variants in the Syrian hamster COVID-19 model

**DOI:** 10.1101/2022.02.22.481491

**Authors:** Kyle Rosenke, Atsushi Okumura, Matthew C. Lewis, Friederike Feldmann, Kimberly Meade-White, W. Forrest Bohler, Amanda Griffin, Rebecca Rosenke, Carl Shaia, Michael A. Jarvis, Heinz Feldmann

**Affiliations:** Laboratory of Virology, National Institute of Allergy and Infectious Diseases, National Institutes of Health; Hamilton, MT, USA; Rocky Mountain Veterinary Branch, National Institute of Allergy and Infectious Diseases, National Institutes of Health; Hamilton, MT, USA; School of Biomedical Sciences, University of Plymouth; Plymouth, Devon, UK; The Vaccine Group Ltd; Plymouth, Devon, UK

## Abstract

The recent emergence of the SARS-CoV-2 Omicron variant of concern (VOC) containing a heavily mutated spike protein capable of escaping preexisting immunity, identifies a continued need for interventional measures. Molnupiravir (MK-4482), an orally administered nucleoside analog, has demonstrated efficacy against earlier SARS-CoV-2 lineages and was recently approved for SARS-CoV-2 infections in high-risk adults. Here we assessed the efficacy of MK-4482 against the earlier Alpha, Beta and Delta VOCs and Omicron in the Syrian hamster COVID-19 model. Omicron replication and associated lung disease in vehicle treated hamsters was reduced compared to the earlier VOCs. MK-4482 treatment inhibited virus replication in the lungs of Alpha, Beta and Delta VOC infected hamsters. Importantly, MK-4482 profoundly inhibited virus replication in the upper and lower respiratory tract of hamsters infected with the Omicron VOC. Consistent with its mutagenic mechanism, MK-4482 treatment had a more pronounced inhibitory effect on infectious virus titers compared to viral RNA genome load. Histopathologic analysis showed that MK-4482 treatment caused a concomitant reduction in the level of lung disease and viral antigen load in infected hamsters across all VOCs examined. Together, our data indicate the potential of MK-4482 as an effective antiviral against known SARS-CoV-2 VOCs, especially Omicron, and likely future SARS-CoV-2 variants.

**One Sentence Summary:** MK-4482 inhibits replication of multiple SARS-CoV-2 variants of concern, including Omicron, in the Syrian hamster COVID-19 model

## INTRODUCTION

Now in its third year, the severe acute respiratory syndrome coronavirus 2 (SARS-CoV-2) coronavirus disease 2019 (COVID-19) pandemic has become characterized by the serial emergence of variants of concern (VOCs) that rapidly and globally replace earlier, previously predominant strains. In November 2021, Omicron (B.1.1.529) emerged to rapidly replace Delta (B.1.617.2), the predominant VOC at the time (*1*). Omicron has shown reduced pathogenicity (*2*) but enhanced transmissibility primarily associated with an increased ability to evade immunity associated with infection by earlier VOCs or through vaccination (*3, 4*). The emergence of such VOCs with immune evasive ability poses a continuous and considerable threat to global control strategies based on present spike-based vaccines, as well as monoclonal antibody-based therapeutics for treatment of more severe COVID-19.

Molnupiravir (MK-4482) is an orally available antiviral nucleoside analogue that targets SARS-CoV-2 polymerase fidelity rather than the spike protein (*5*). It has been authorized for emergency use against SARS-CoV-2 in high-risk adults (*6, 7*). Results from recent in vitro studies suggest that molnupiravir retains activity against multiple VOCs including Omicron (*8*). However, experience with therapeutics targeting SARS-CoV-2 (*9*), as well as other emerging viruses (*10*) have shown a frequent disconnect between efficacy observed *in vitro* compared to therapeutic *in vivo* effect. In the present study we investigate the ability of MK-4482 to inhibit several SARS-CoV-2 VOCs (Alpha, Beta, Delta and Omicron) in the Syrian hamster COVID-19 model.

## RESULTS

### MK-4482 treatment inhibits replication of multiple VOCs, including Omicron

Syrian hamsters were randomly divided into vehicle-or MK-4482-treatment groups, and infected intranasally with 10^3^ TCID_50_ as previously established (*11*) to test in vivo efficacy of MK-4482 against the SARS-CoV-2 Alpha, Beta, Delta and Omicron VOCs. Treatment using 250mg/kg MK-4482, administered by oral gavage, was initiated 12h after infection. Treatment was continued every 12h for the established total daily dose of 500mg/kg (*12*). Hamsters were monitored daily for clinical signs and oral swabs were collected 2 and 4 days post-infection (dpi). At 4 dpi hamsters were euthanized and tissues collected for viral RNA load and infectious titer determination (Fig. 1A).

**Fig 1.**
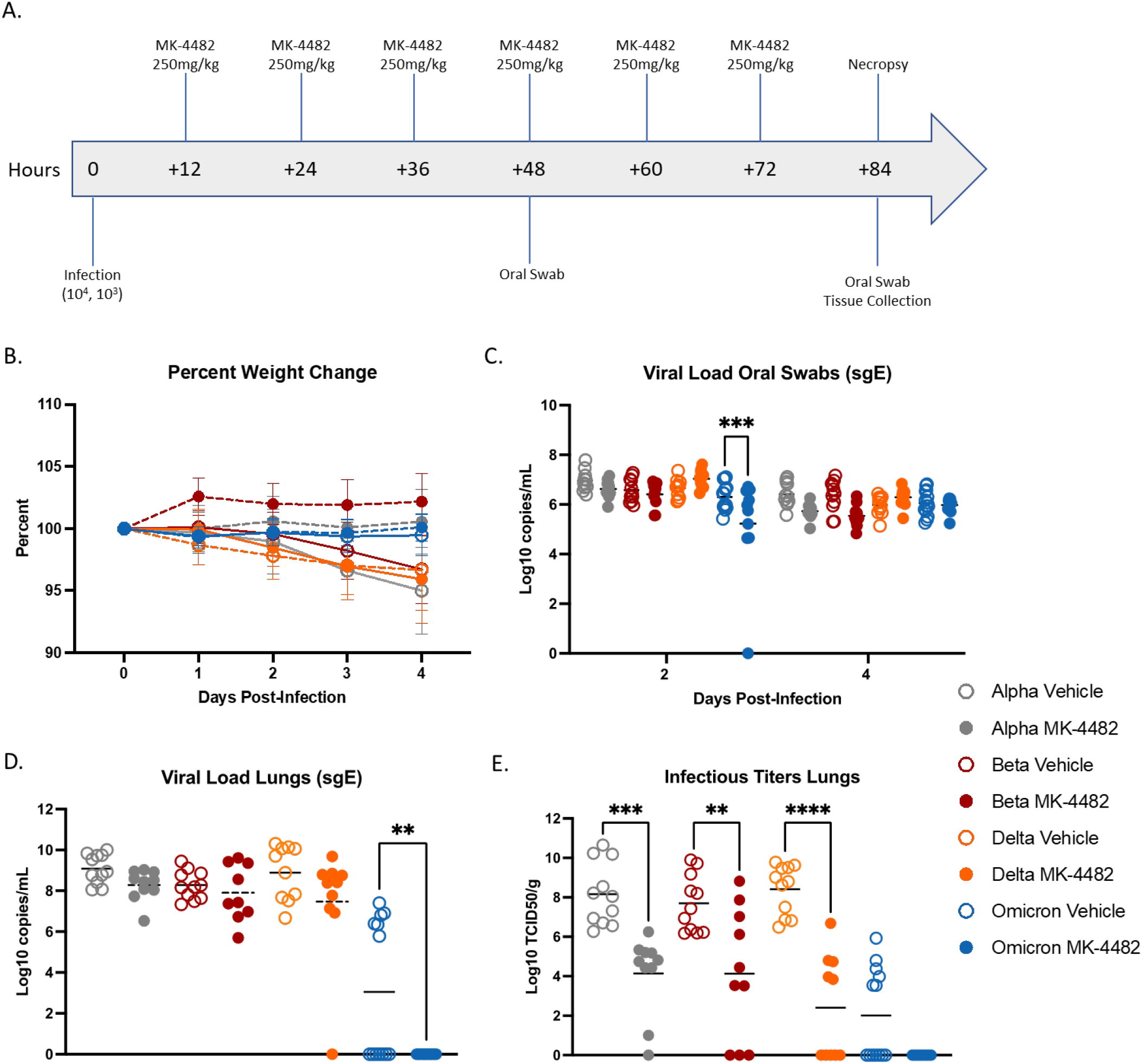
MK-4482 efficacy on upper and lower respiratory tract viral load and infectivity in hamsters infected with Alpha, Beta, Delta and Omicron SARS-CoV-2 VOCs. **(A) Experimental design.** Hamsters (N =11 vehicle, 10 treatment) were intranasally infected with 10^3^ TCID_50_ of the different SARS-CoV-2 VOCs. Treatment was started 12 hours post infection and continued every 12 hours. Oral swabs were collected on 2 and 4 dpi and animals were necropsied on 4 dpi for tissue collection. **(B) Clinical presentation**. Changes in body weight were recorded over the entire study period of 4 days. **(C) Viral RNA load in oral swabs**. Alpha, Beta, Delta and Omicron RNA loads were determined by quantitative RT-PCR targeting sgE RNA as a surrogate for replication and shedding. **(D) Viral RNA loads in lung tissue**. Alpha, Beta, Delta and Omicron RNA loads were determined by quantitative RT-PCR targeting sgE as a surrogate for replication. **(E) Infectious viral titers in lung tissue**. Alpha, Beta, Delta and Omicron infectivity was determined using a tissue culture infectious dose (TCID) assay and are presented as TCID_50_/gram tissue. Statistical differences in viral load and infectious virus titers in each study arm were assessed by ordinary one-way ANOVA.

Hamsters in vehicle and MK-4482 treatment groups remained largely asymptomatic with noticeable but not significant weight loss (3-4%) in vehicle treated animals over the 4-day study period (Fig. 1B). Quantitative RT-PCR (qRT-PCR) targeting subgenomic viral E gene RNA (sgE) in oral swabs and lung tissue was used to quantify replication in the upper and lower respiratory tract, respectively. sgE loads in oral swabs from vehicle treated animals on 2 and 4 dpi ranged from 5 to 8 log_10_ copies/mL confirming infection by all VOCs with the exception of a single Omicron animal (Fig. 1C). MK-4482 treatment resulted in lower replication in oral swabs for all VOCs except Delta, a result that was only statistically significant for Omicron at 2 dpi (Fig. 1C). sgE loads in lung tissue ranged from 7 to 10 log_10_ copies/g for Alpha, Beta and Delta in vehicle treated groups, but was notably lower (0-7 log_10_ copies/g) for Omicron (Fig. 1D). MK-4482 treatment resulted in a reduction in sgE lung loads for all VOCs and was below the limit of detection for Omicron, a result that was statistically significant (Fig. 1D). Similarly, infectious virus titers in lung tissue ranged from 6-11 log_10_ TCID_50_/g for Alpha, Beta and Delta vehicle treated groups that was significantly lower for Omicron (0 to 6 log_10_ TCID_50_/g) (Fig. 1E). Infectious titers were substantially more reduced by MK-4482 than sgE loads. This was observed for all VOCs, with no infectious virus detected in lung tissue from any Omicron infected hamster (Fig. 1E).

### MK-4482 treatment reduces pathology and virus antigen in the lung for multiple VOCs, including Omicron

Histopathologic analysis of lung tissues revealed minimal to moderate broncho-interstitial pneumonia for vehicle treated Alpha, Beta and Delta VOC infected hamsters. Omicron infected animals showed only mild lesions reflecting reduced virus replication of this VOC in the lower respiratory tract (*13*) (Fig. 2). In general lesions were as previously described (*11*). They consisted of bronchiolar and alveolar inflammation most prominent around terminal bronchioles with neutrophils, macrophages, necrotic debris, fibrin and edema in associated alveolar spaces as well as variable amounts of septal inflammation and thickening (fig. S1). Other features included single and small clusters of necrotic respiratory epithelium in large and medium caliber bronchioles and vasculitis in locally associated blood vessels (fig. S1). Lesions were reduced in all MK-4482 treated hamsters as noticeably shown on the lung hematoxylin and eosin (H&E) overview stain (Fig. 2) and the more detailed magnified H&E (fig. S1). Immunohistochemical labeling of affected and non-affected bronchioles and alveoli was common throughout, but especially at the margins of regions of pneumonia for vehicle treated hamsters infected with Alpha, Beta and Delta VOCs. Labeling was only infrequently found with hamsters infected with the Omicron VOC (Fig 2; fig. S1)). Immunoreactivity was consistently less frequent in MK-4482 treated hamsters for each group with the least frequent immunoreactivity in the treated Omicron infected hamsters (Fig. 2; fig. S1)

**Fig 2.**
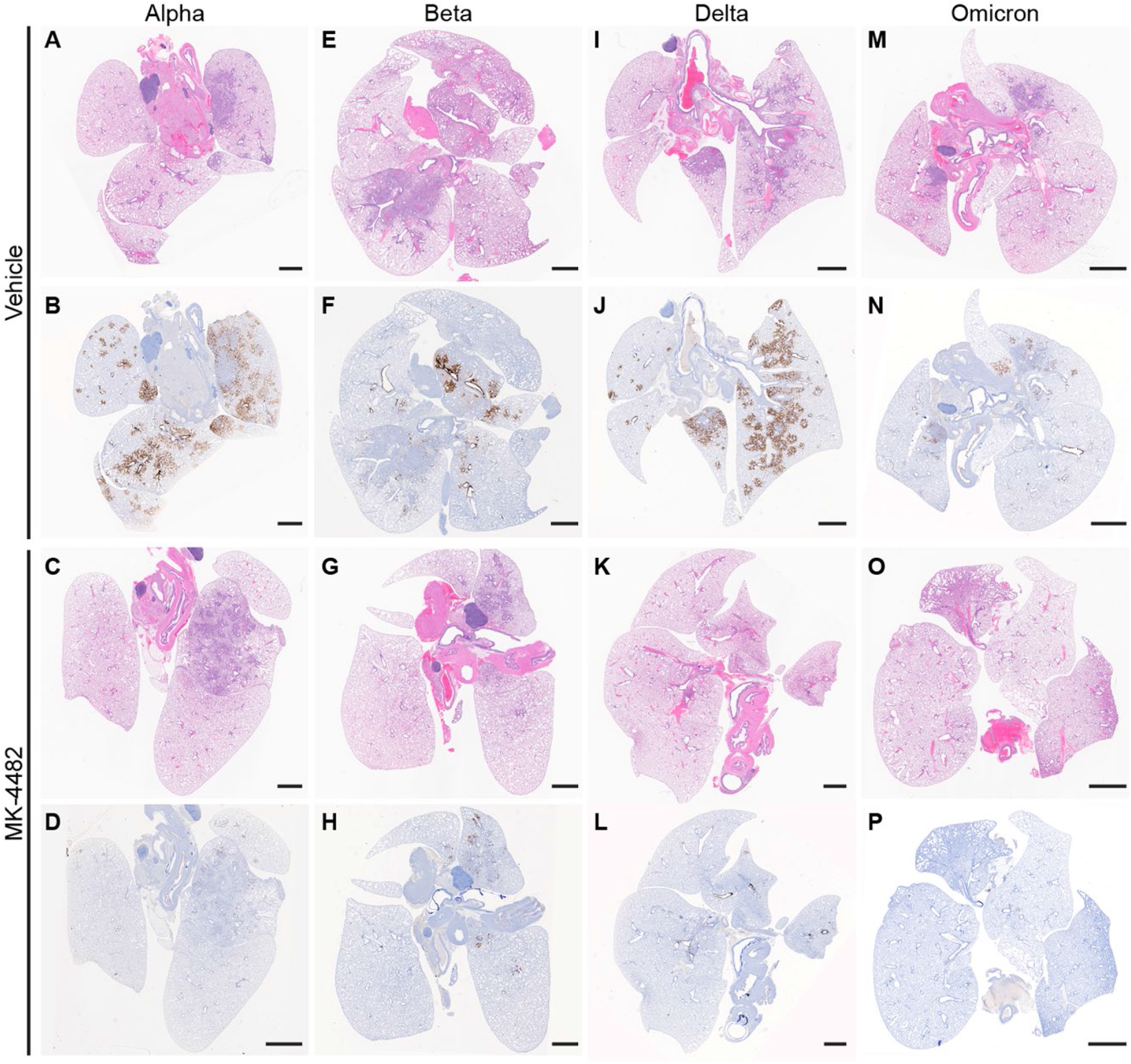
MK-4482 efficacy on lung tissue from Alpha, Beta, Delta and Omicron infected hamsters. The experimental design is shown in Figure 1A. Lung tissue was collected on 4 dpi and prepared for hematoxylin and eosin (H&E) stain and immunohistochemistry (IHC) using an antibody directed against the SARS-CoV-2 nucleocapsid protein. (**A, E, I, M**.) **Lung histopathology**. H&E stain of lung tissue from vehicle treated hamsters infected with Alpha, Beta, Delta and Omicron VOC, respectively. Areas of consolidation representing foci of broncho-interstitial pneumonia. (**B, F, J, N**) **Lung immunohistochemistry**. SARS-CoV-2 nucleoprotein detection in lung section from vehicle-treated hamsters infected with Alpha, Beta, Delta and Omicron, respectively. Corresponding foci of immunoreactivity (brown color) are shown. (**C, G, K, O**) **Lung histopathology**. H&E stain of lung tissue from MK-4482 treated hamsters infected with Alpha, Beta, Delta and Omicron VOC, respectively. Areas of consolidation representing foci of broncho-interstitial pneumonia. (**D, H, L, P) Lung immunohistochemistry**. SARS-CoV-2 nucleoprotein detection in lung section from MK-4482 treated hamsters infected with Alpha, Beta, Delta and Omicron, respectively. Corresponding foci of immunoreactivity (brown color) are shown. Bars = 500 μm.

### Inhibitory effect of MK-4482 on Omicron VOC resists higher virus challenge

We repeated the MK-4482 treatment study with a higher Omicron challenge dose (10^4^ TCID_50_) to account for the decreased replication of the Omicron VOC in lung tissue (*13*). The study design remained the same as described above (Fig. 1A), but trachea tissue was collected at 4 dpi as an additional target tissue. The higher Omicron challenge dose did not increase clinical disease severity and weight loss remained similar to animals challenged with the lower 10^3^ TCID_50_ dose (Figs. 1B,3A). At 2 and 4 dpi, oral swab sgE loads from vehicle treated animals ranged from 5 to 7 log_10_ copies/mL, which were lower in MK-4482 treated animals (<1 log_10_ copies/mL), a difference that was not statistically significant (Fig. 3B). Infectious virus in oral swabs from vehicle treated animals ranged from 4 to 7 log_10_ TCID_50_/mL (Fig. 3C), which was significantly reduced by several log_10_ following MK-4482 treatment (Fig. 3C). sgE loads and infectious titers were also determined in trachea and lung tissues at 4 dpi. In contrast to trachea, approximately half the animals in the vehicle treated group lacked detectable virus in the lung (by TCID_50_) consistent with lower replication of Omicron in this tissue in general (*13*). Remarkably, MK-4482 treatment reduced sgE loads and infectious virus to below the limit of detection in lung tissue of all animals, corresponding to an average reduction by >4 log_10_ (Fig. 3D) and >2 log_10_ (Fig. 3E), respectively. Histopathologic analysis of the lungs confirmed the findings showing reduced lesions and no viral antigen in the MK-4482 treated Omicron infected hamsters (fig. S2). Interestingly, sgE loads were several log_10_ higher in trachea (9 to 11 log_10_ copies/g) compared to lung tissue of vehicle treated hamsters. sgE loads were less than a log_10_ lower in trachea tissue from MK-4482 treated animals (Fig. 3D). However, infectious virus in trachea tissue dropped precipitously from a median of 5 log_10_ TCID_50_/g in vehicle treated animals to below the limit of detection in MK-4482 animals (Fig. 3D,E).

**Fig 3.**
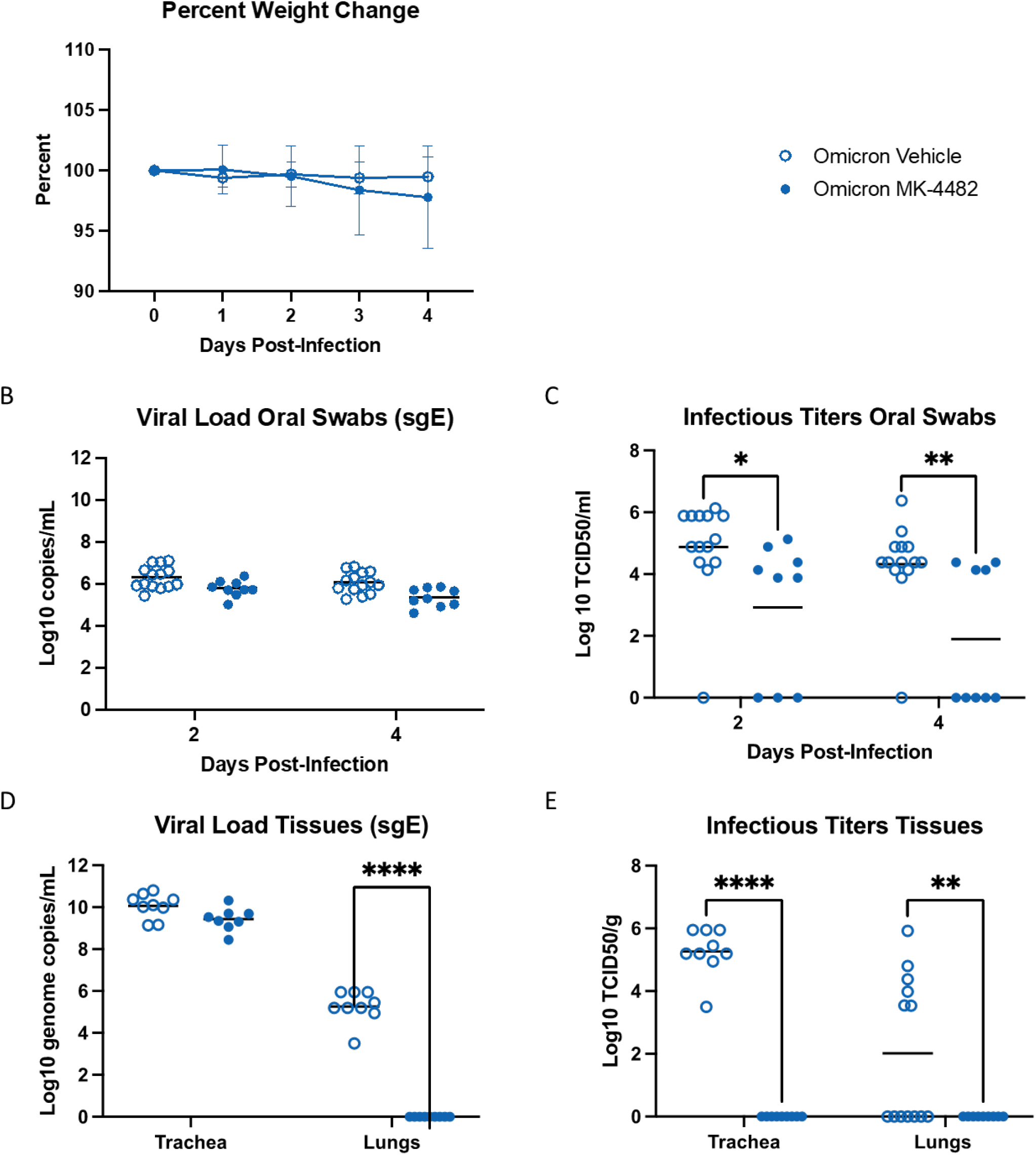
MK-4482 efficacy on upper and lower respiratory tract viral load and infectivity in hamsters infected with high dose Omicron SARS-CoV-2 VOC. The experimental design is shown in Figure 1A. For this experiment we used an Omicron challenge dose of 10^4^ TCID_50_ and collected trachea as an additional tissue. **(A) Clinical presentation**. Changes in body weight were recorded for the entire study period of 4 days. **(B) Omicron RNA load in oral swabs**. Viral loads were determined by quantitative RT-PCR targeting sgE RNA as a surrogate for replication and shedding. **(C) Infectious Omicron titers in oral swabs**. Viral infectivity was determined using a TCID_50_ assay and are presented as TCID_50_/gram tissue. **(D**) **Omicron loads in trachea and lung tissues**. Viral loads were determined by quantitative RT-PCR targeting sgE RNA as a surrogate for replication. **(E) Infectious Omicron titers in trachea and lung tissues**. Viral infectivity was determined using a TCID_50_ assay and are presented as TCID_50_/gram tissue. Statistical analysis was performed in Prism 9. Statistical differences in viral load and infectious virus titers in each study arm were assessed by ordinary one-way ANOVA.

## DISCUSSION

Since SARS-CoV-2 emerged in late 2019, there have been at least 13 variants of interest (VOI) or VOCs circulating throughout the world (*14*). As each variant is likely to become more adapted at spread and avoiding preexisting immunity developed by previous infection or vaccination, refining existing and developing new treatment options becomes critically important for our response to the SARS-CoV-2 pandemic. In this study we continue our evaluation of MK-4482 against SARS-CoV-2 infection, testing its efficacy against several SARS-CoV-2 VOCs in the Syrian hamster model. MK-4482 was efficacious against all VOCs with significant reduction of virus replication in the lower respiratory tract. This was particularly evident for the Omicron VOC with absence of any infectious virus in trachea and lung samples. Thus, molnupiravir remains potent against SARS-CoV-2 variants including Omicron, likely positively affecting COVID-19 patient outcome.

Important for transmission, a key driver of any epidemic and pandemic, is the impact of an antiviral intervention on virus shedding. In contrast to SARS-CoV-2 replication in the lower respiratory tract, MK-4482 treatment only slightly decreased replication and shedding from the upper respiratory tract for the Alpha, Beta and Delta VOCs. Interestingly, MK-4482 treatment significantly reduced replication and shedding of Omicron, indicating potent efficacy against the VOC driving the current SARS-CoV-2 pandemic. This is a somewhat surprising but rather encouraging result with the potential of positively influencing the course of the current pandemic wave.

The antiviral activity of MK-4482 is not affected by mutations in the spike protein and remains active against all variants of SARS-CoV-2. Thus, MK-4482 remains a potent alternative to monoclonal antibody treatment which is rather vulnerable to mutations in the spike protein (*8*). Although Omicron already has mutations in the viral polymerase protein (*15*), the target of MK-4482 antiviral action, the drug remains effective against this VOC with significant reduction of viral replication in the upper and lower respiratory tract. It remains to be seen whether future variants that may develop under more widespread clinical use of molnupiravir will acquire further mutations in the polymerase resulting in reduced drug activity or even loss of drug efficacy. Nevertheless, serial in vitro passage studies of several RNA viruses (*16, 17*) as well as related coronaviruses (*18*) under suboptimal levels of the active metabolite of MK-4482 indicate a high genetic barrier of the drug to the acquisition of viral resistance. Selective and targeted clinical use of the drug as well as the consideration of combination therapy with synergistic antivirals or other therapeutic approaches (*19*) may also help to reduce the development of such drug resistance.

We found a marked discrepancy between viral sgRNA loads and infectious titers for all VOCs. Given the mechanism of action of MK-4482 (*5*) this may not be surprising as mutated, replication incompetent viral RNA will still be detected by qRT-PCR analysis but not by infectivity assays. This observation emphasizes the need to consider the specific method of analysis being used to assess antiviral activity, as viral sgRNA loads are often used as the single measure for SARS-CoV-2 replication. Our results strongly suggest that infectious titers are critical for a complete investigation and required to determine the efficacy of antiviral drugs and likely also vaccines. Histopathology serves as an important confirmation for virus replication and provides helpful pathologic images.

Our study has limitations. The hamster model does not accurately represent COVID-19 disease for all human age- and comorbidity-associated subpopulations as the animals only develop mild-to-moderate disease (*11, 20, 21*), but this is a drawback of most current animal models (*22*). The more recently identified dwarf hamster model is associated with higher disease severity, but the extremely acute disease progression with animals reaching clinical endpoints within as quickly as 3 dpi (*23*) does not parallel the more prolonged course of severe disease in humans (*24*). In all cases, the current intranasal delivery of the challenge virus, whilst mimicking the most biologically relevant route of infection in humans, may interfere with readout parameters for the upper respiratory tract negatively impacting analysis of virus replication and shedding. Future transmission studies need to confirm the impact of MK-4482 on virus shedding and ultimately transmission to naïve animals. A study awaiting peer-review has shown an inhibitory effect of MK-4482 on shedding and transmission in the upper respiratory disease ferret model (*25*), which is encouraging; transmission studies are more difficult to perform in the hamster (*26*) and should be targeted for future experiments. The greatest limitation in the current situation is the incomplete experience with molnupiravir in clinical studies against the Omicron VOC. Those studies are underway, and results are expected in the near future. Nevertheless, in vitro studies (*8*), preclinical data from animal models such as our study, and previous clinical trials against earlier SARS-CoV-2 VOCs (*6*) strongly support the use of molnupiravir for mild-to-moderate COVID cases of Omicron in high-risk adults as well as possibly for postexposure prophylaxis.

In conclusion, our data demonstrate potent efficacy of MK-4482 against the dominant Omicron VOC in a key well-established in vivo COVID-19 disease model. We also demonstrated broad efficacy of MK-4482 against earlier globally distributed VOCs identifying molnupiravir as an important intervention for existing and likely future VOCs. The potent efficacy and favorable route of oral administration support further clinical development and broader use of molnupiravir as a key treatment option for the SARS-CoV-2 pandemic.

## MATERIALS AND METHODS

### Biosafety and ethics

SARS-CoV-2 work was approved by the Institutional Biosafety Committee (IBC) and performed in high biocontainment at Rocky Mountain Laboratories (RML), NIAID, NIH. IBC-approved Standard Operating Protocols were followed for sample removal from biocontainment. Institutional Animal Care and Use Committee approved all animal work which was performed by certified staff in an Association for Assessment and Accreditation of Laboratory Animal Care International accredited facility. The institution’s guidelines for animal use, the guidelines and basic principles in the NIH Guide for the Care and Use of Laboratory Animals, the Animal Welfare Act, United States Department of Agriculture and the United States Public Health Service Policy on Humane Care and Use of Laboratory Animals were followed. Syrian hamsters were group housed in HEPA-filtered cage systems enriched with nesting material and were provided with commercial chow and water *ad libitum*. Animals were monitored at least twice daily.

### Syrian hamster study design

Male and female 8–10-week-old hamsters were divided into vehicle (N=15 for Omicron infected, N=11 Alpha, Beta, Delta infected) or treatment (N=10 for Alpha, Beta, Delta, Omicron infected; N=9 for the Omicron high dose infected) groups prior to infection and treatments with MK-4482 (MedChemExpress). MK-4482 was dissolved in DMSO and then resuspended in sterile saline for delivery at 250mg/kg. Hamsters were treated 12 hours following infection, and treatment was continued every 12 hours until the completion of the study 84 hours post-infection (day 4). Vehicle control animals received the same dosing schedule and volume as VOC infection groups. All groups were infected intranasally with 10^3^ or 10^4^ TCID_50_ (high dose) of SARS-CoV-2 (25 µL/nare). Animal weights were collected once daily, and animals were monitored twice daily for disease signs and progression. All procedures were performed on anesthetized animals. Oral swabs were collected on days 2 and 4 post-infection. Animals were euthanized on day 4 post-infection and trachea and lung tissues were collected at necropsy for analysis.

### Virus and cells

The SARS-CoV-2_2hCOV_19_England_204820464_2020 isolate (B.1.1.7; Alpha) was provided by BEI Resources. The stock was sequence confirmed and had amino acid changes at ORF1AB (D3725G: 13%) and ORF1AB (L3826F: 18%) when aligned to the GISAID sequence (GISAID # EPI_ISL_683466). SARS-CoV-2 isolate nCoV-hCoV-19/USA/MD-HP01542/2021 (B.1.351, Beta) was provided by Andy Pekosz (Johns Hopkins). The virus stock was sequence confirmed and found to have amino acid changes at NSP5 (P252L: 17%) and NSP6 (L257F: 57%) when aligned to the GISAID sequence (GISAID # EPI_ISL_890360). SARS-CoV-2 variant hCoV-19/USA/KY-CDC-2-4242084/2021 (B.1.617.2, Delta) was obtained with contributions from B. Zhou, N. Thornburg and S. Tong (Centers for Disease Control and Prevention, USA). SARS-CoV-2 variant hCoV-19/USA/GA-EHC-2811C/2021 (B.1.1.529, Omicron, EPI_ISL_7171744) was obtained from Mehul Suthar, Emory University. All viral stocks were sequenced via Illumina-based deep sequencing to confirm identity and exclude any contaminants. Virus propagation was performed in DMEM (Sigma) supplemented with 2% fetal bovine serum (Gibco), 1 mM L-glutamine (Gibco), 50 U/ml penicillin and 50 μg/ml streptomycin (Gibco). Vero E6 cells, kindly provided by R. Baric, University of North Carolina, were maintained in DMEM (Sigma) supplemented with 10% fetal calf serum, 1 mM L-glutamine, 50 U/mL penicillin and 50 μg/mL streptomycin.

### Viral genome detection

qPCR was performed on RNA extracted from swabs or tissues (30 mg or less) using QiaAmp Viral RNA kit. A one-step real-time RT-PCR assay was used to amplify a portion of the E gene to detect subgenomic RNA (*27*). Dilutions of RNA standards counted by droplet digital PCR were run in parallel and used to calculate viral RNA genome copies. The Rotor-Gene probe kit (Qiagen) was used to run the PCRs according to the instructions of the manufacturer.

### Virus titration assay

Virus end-point titrations were performed in Vero E6 cells. Briefly, tissue was homogenized in 1ml DMEM using a TissueLyzer (Qiagen) and clarified by low-speed centrifugation. Cells were inoculated with 10-fold serial dilutions of homogenized lung samples or oral swabs in 100 µl DMEM (Sigma-Aldrich) supplemented with 2% fetal bovine serum, 1 mM L-glutamine, 50 U/ml penicillin and 50 µg/ml streptomycin. Cells were incubated for six days and then scored for cytopathogenic effects (CPE) and TCID_50_ was calculated via the Reed-Muench formula.

### Histopathology

Tissues were embedded in Pureaffin paraffin polymer (Cancer Diagnostics, Durham, NC, USA) and sectioned at 5 µm for hematoxylin and eosin (H&E) staining. For immunohistochemistry (IHC), tissues were processed using the Discovery Ultra automated stainer (Ventana Medical Systems) with a ChromoMap DAB kit (Roche Tissue Diagnostics cat#760-159). Specific immunoreactivity was detected using the GenScript U864YFA140-4/CB2093 NP-1 SARS-CoV-2-specific antiserum at a 1:1000 dilution. The secondary antibody was an anti-rabbit IgG polymer (cat# MP-6401) from Vector Laboratories ImPress VR. As all animals in this study were inoculated with SARS-CoV-2, uninfected tissues from previous age-matched porcine studies were used for control tissues.

### Statistical analyses

Statistical analysis was performed in Prism 9. The difference in weight, viral load and infectious titers between study arms was assessed by ordinary one-way ANOVA.

## LIST OF SUPPLEMENTARY MATERIAL

**Fig. S1 to Fig. S2**

## ACKNOWLEDGMENTS

The authors thank staff members of the Laboratory of Virology and the Genomics Unit of the Research Technology Branch (both National Institutes of Allergy and Infectious Diseases (NIAID) and National Institutes of Health (NIH) for their efforts in providing workable SARS-CoV-2 stocks. The authors also wish to thank the Rocky Mountain Veterinary Branch (NIAID, NIH) for animal care and husbandry. The authors are thankful to Anita Mora, Visual and Medical Arts Unit, NIAID, NIH) for help with graphical design. Isolate SARS-CoV-2_2hCOV_19_England_204820464_2020 isolate (B.1.1.7; Alpha) was obtained through BEI Resources (Bassam Hallis, Sujatha Rashid), NIAID (Ranjan Mukul, Kimberly Stemple) and the NIH. Isolate hCoV-19/USA/MD-HP01542/2021 was obtained from Andrew Pekosz, John Hopkins Bloomberg School of Public Health. Isolate hCoV-19/USA/GA-EHC-2811C/2021 was obtained from Mehul Suthar, University Emory School of Medicine. The following reagent was deposited by the Centers for Disease Control and Prevention and obtained through BEI Resources, NIAID, NIH: hCoV-19/USA/KY-CDC-2-4242084/2021. Vero E6 cells were obtained from Dr. Ralph Baric at the University of North Carolina.

## Funding

The work was mainly funded by the Intramural Research Program of NIAID, NIH and through the Vaccine Group Ltd and the University of Plymouth.

## Author contributions

KR, MAJ and HF contributed to the design, execution and data analysis, and writing of the manuscript. FF, KMW, AO, AG, WFB and ML contributed experiment support and data analysis. HF and MAJ secured funding of the study. All authors reviewed and contributed to preparation of the final manuscript.

## Competing interests

MAJ is employed, in part, by the Vaccine Group Ltd. All authors declare that they have no competing interests.

## Data and materials availability

All data is presented here or in Supplementary Material. For additional information can be requested through the corresponding authors.

## Disclaimer

The opinions, conclusions and recommendations in this report are those of the authors and do not necessarily represent the official positions of the National Institute of Allergy and Infectious Diseases (NIAID) at the National Institutes of Health (NIH), The Vaccine Group Ltd or the University of Plymouth.

## SUPPLEMENTARY MATERIAL

**Fig. S1.**
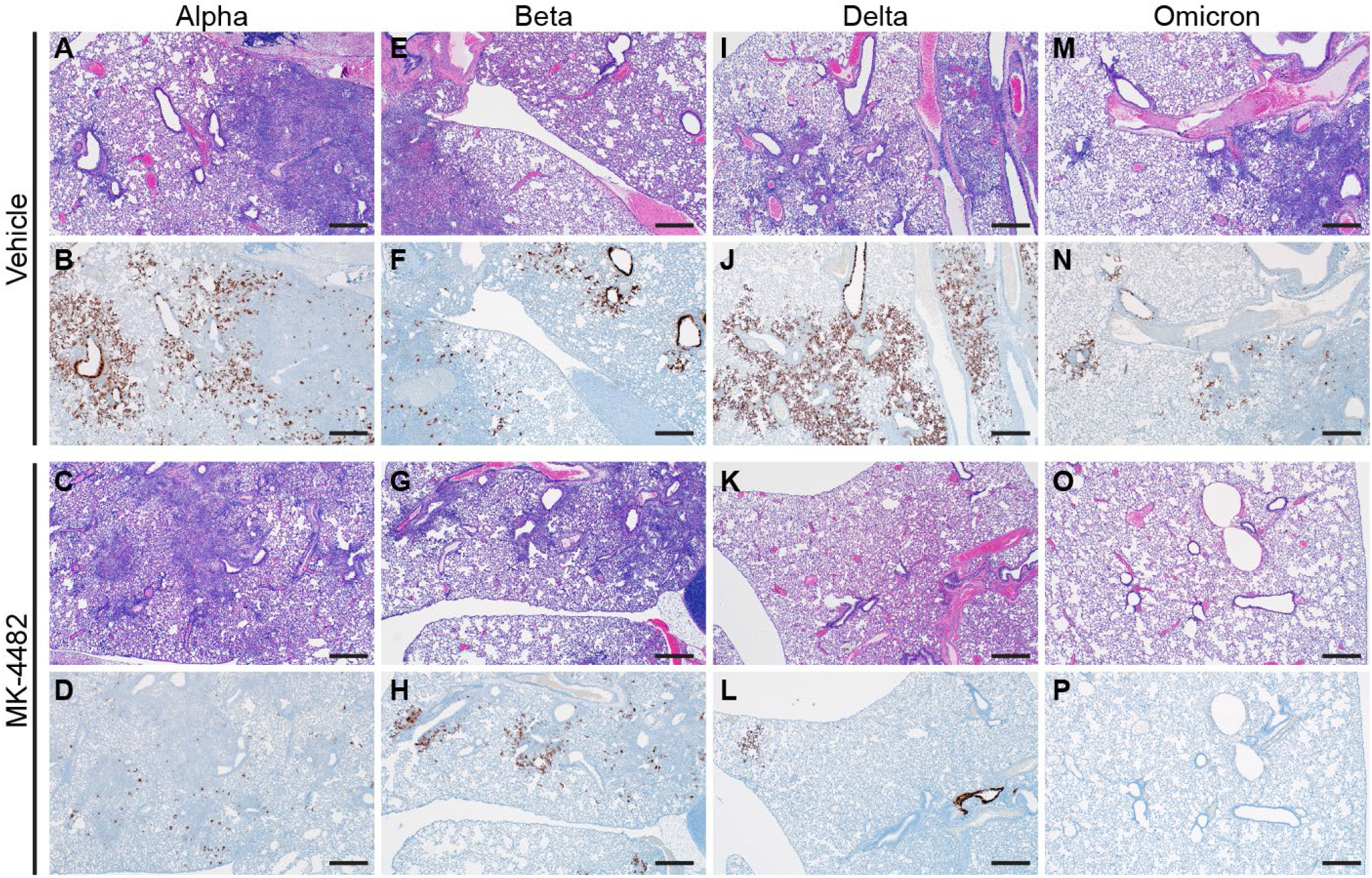
MK-4482 efficacy on lung pathology in hamsters infected with multiple SARS-CoV-2 VOCs. The experimental design is shown in Figure 1A. Lung tissue was collected on 4 dpi and prepared for hematoxylin and eosin (H&E) stain and immunohistochemistry (IHC) using an antibody directed against the SARS-CoV-2 nucleocapsid protein. **(A, E, I, M) Lung H&E (40x)**. H&E stain from vehicle treated hamsters infected with Alpha, Beta, Delta and Omicron VOC, respectively, showing bronchointerstitial pneumonia and vasculitis. **(B, F, J, N) Lung IHC (40x)**. SARS-CoV-2 nucleoprotein detection in lung section from vehicle-treated hamsters infected with Alpha, Beta, Delta and Omicron, respectively, exhibiting immunoreactivity associated with areas of pneumonia (brown color). **(C, G, K, O) Lung H&E (40x)**. H&E stain from MK-4482 treated hamsters infected with Alpha, Beta, Delta and Omicron VOC, respectively, showing reduced bronchointerstitial pneumonia. **(D, H, L, P) Lung IHC (40x)**. SARS-CoV-2 nucleoprotein detection in lung section from MK-4482 treated hamsters infected with Alpha, Beta, Delta and Omicron VOC, respectively, exhibiting reduced or no immunoreactivity. Each panel shows a lung section from a representative hamster. Bar = 500 μm

**Fig. S2.**
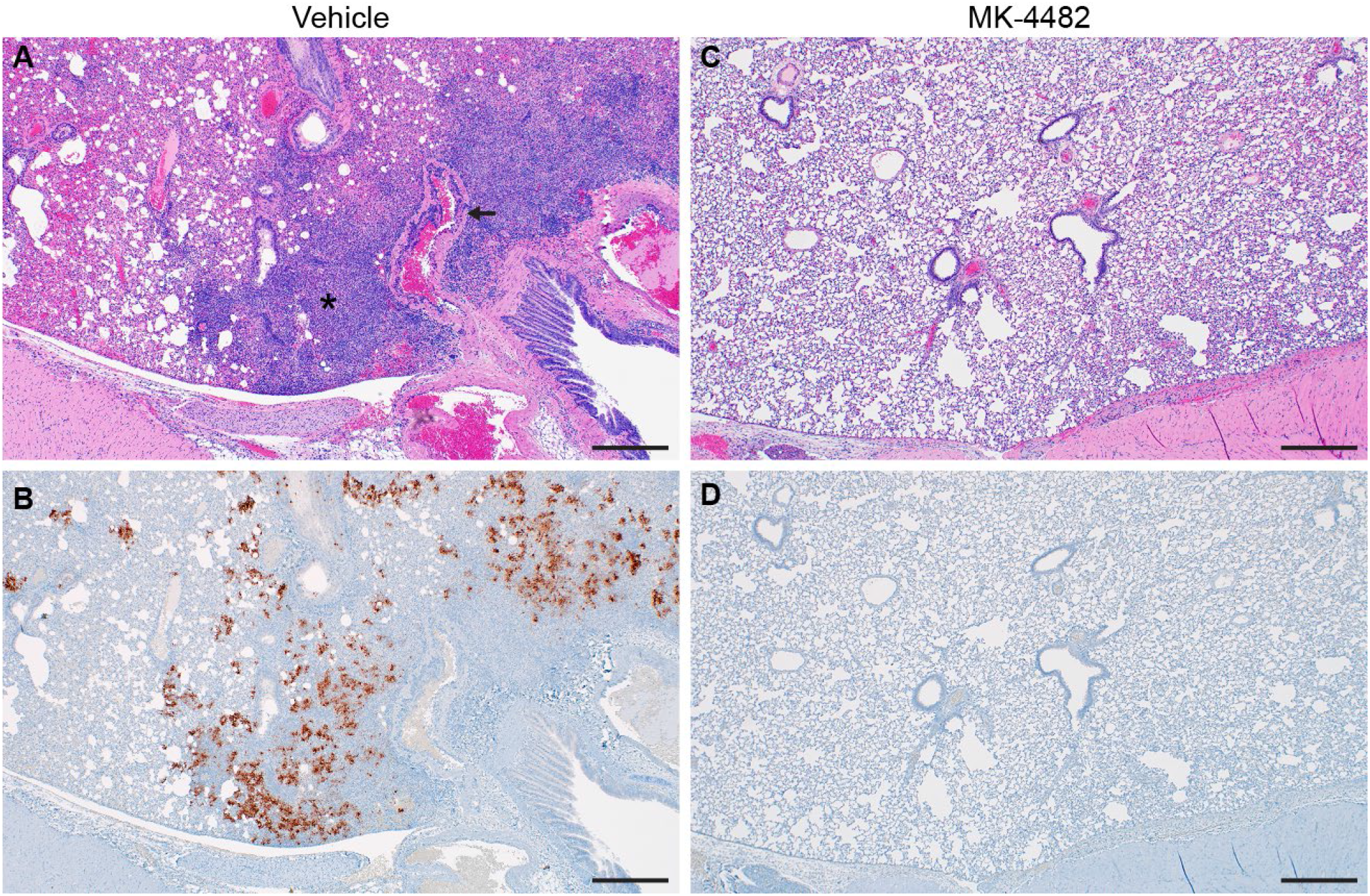
MK-4482 efficacy on lung pathology in hamsters infected with a high dose of Omicron SARS-CoV-2 VOC. The experimental design is shown in Figure 1A. Lung tissue was collected on 4 dpi and prepared for H&E stain and IHC using an antibody directed against the SARS-CoV-2 nucleocapsid protein. **(A) Lung H&E (40x)**. Lung H&E of a vehicle treated hamster infected with Omicron exhibiting a focus of bronchointerstitial pneumonia (asterisk) and vasculitis (arrow). **(B) Lung IHC (40x)**. Lung IHC from a vehicle treated hamster infected with Omicron exhibiting frequent immunoreactivity associated with focus of pneumonia (brown color). **(C) Lung H&E (40x)**. Lung H&E of a MK-4482 treated hamster infected with Omicron showing normal pathology. **(D) Lung IHC (40x)**. Lung IHC of a MK-4482 treated hamster infected with Omicron exhibiting absence of immunoreactivity. Each panel shows a lung section from a representative hamster. Bar = 500 μm

